# From planktonic to sedentary lifestyle: Molecular dissection of the establishment and maintenance of mycobacterial biofilm

**DOI:** 10.64898/2026.07.04.736460

**Authors:** Himalaya Naik, Rohit Satardekar, Raju Mukherjee, Vikas Jain

## Abstract

Biofilm represents a complex aggregation of bacteria embedded within a self-produced extracellular polymeric substance (EPS). We investigated the characteristics of mycobacterial biofilm using *Mycobacterium smegmatis* (Msm) as model organism. By combining transcriptomic (RNA-seq) and proteomic (LC-MS) analyses, the research captures dynamic changes during the establishment and maturation of the biofilm. Transcriptomics analysis showed a distinct gene expression profile as compared to its planktonic form. Interestingly, clear differences were seen between initial (∼2-day old) and mature (∼5-day old) biofilm stages, highlighting phasic gene expression throughout biofilm development. Marked alteration in oxidative stress-related genes and energy metabolism from ATP to NADH was observed. Furthermore, quantitative mass spectrometry-based proteome examination of EPS showed an abundance of cytoplasmic proteins present differentially between initial and mature biofilm stages. Pathway enrichment revealed enhanced oxidative stress responses and metabolic shifts in mature biofilms, including upregulation of NADH dehydrogenase and downregulation of ATP synthase, indicating altered energy metabolism. Our findings thus provide insights into the molecular adaptations, including production of mycofactocin, occurring during mycobacterial biofilm establishment and maturation, and advance our understanding of mycobacterial biofilm physiology.

## Introduction

Biofilm is a complex multicellular structure glued with extracellular polymeric substances (EPS), harnessed by bacteria in response to external stress, including host response and antibiotics.^1^ Bacterial cells residing in the biofilms are metabolically active, drug-tolerant, and show persistence compared to the planktonic counterparts.^2^ It has been hypothesized that the unresponsiveness and the emergence of resistance towards antimicrobial drugs during pathogenic infection are mainly due to the formation of biofilms.^3^ EPS provides mechanical stability and acts as a barrier against antibiotic treatment.^4,5^ This complex multicellular behavior of bacteria in biofilm is orchestrated by proteins, one of the major components in the EPS.^6^

Mycobacteria contain various infectious species including *Mycobacterium tuberculosis* (Mtb) and other Non-Tuberculous Mycobacteria. They show a high propensity to form pellicular biofilms and therefore are cultured in the presence of detergents for proper planktonic growth.^7^ Previous research on *Mycobacterium smegmatis (Msm)* biofilms has highlighted the significance of various factors, including the role of surface free lipids, iron homeostasis, and short chain mycolic acids in biofilm establishment. Several mutants have been identified lacking the ability to form biofilms or having an altered biofilm phenotype.^8–10^ Abrogation of glycopeptidolipid biosynthesis leads to disruption in biofilm formation. *Msm* biofilms are greatly affected by iron availability, and the biofilm shows increased expression of siderophore genes.^9^ Both *Mtb* and *Msm* show accelerated biofilm formation in the presence of DTT, which, however, is BSA-dependent, indicating the role of extracellular polymeric substrates.^11^ This indicates the importance of proteins in EPS.

Studies in other bacteria, such as *Pseudomonas aeruginosa*, revealed distinct physiological states during biofilm development, with at least five different stages identified through temporal transcriptomic profiling.^12^ This study showed that a large number of proteins were differentially produced during various stages of biofilm development, with maximum changes observed in the cells present in the mature biofilm as compared to those present as planktonic. In *Escherichia coli*, transcriptomic analysis of biofilms revealed differential expression of over 600 genes, with 9% of the genome being activated and 4.5% repressed in the cells in biofilm compared to in planktonic form.^13^ Similarly, in *Bacillus subtilis*, transcriptomic analysis identified 519 genes as differentially expressed during biofilm formation, with more than 55% of these expressed at only one of the three time points examined, indicating temporal control of gene expression.^14^ Proteomic studies in *B. cereus* and *Burkholderia pseudomallei* also showed distinct and reproducible protein patterns between biofilms of different ages.^15,16^ These findings highlight the importance of studying the temporal changes of biofilm formation. It also demonstrates that biofilm development involves complex, stage-specific changes in gene expression and protein production and secretion to form the extracellular matrix. Given the importance of biofilm in infection and persistence, processes and components specific for different stages could serve as targets for intervention. While transcriptomics studies analyzing the biofilm have been conducted in mycobacteria,^17–19^ there is a lack of such stage-specific, temporal, and transitional analysis for mycobacterial biofilms.

In this study, using a multi-omics approach with transcriptomics and proteomics, we examined mycobacterial biofilm at different stages. To understand how mycobacteria form biofilm and, more importantly, stay in this state for a long duration, we examined transcriptome and proteome profiles of the biofilm of a model mycobacteria, *M. smegmatis*. We here present a detailed analysis of the factors responsible for biofilm formation as well as its long-term maintenance. Proteome analysis of EPS revealed the abundance of cytoplasmic proteins in the EPS, potentially indicating their role in biofilm nucleation. We also show a shift in energy metabolism from ATP to NADH/NAD+ along with the importance of electron acceptors during the transition from planktonic to biofilm. Our study thus provides a global view of the transcriptional and proteomic adaptations involved in biofilm establishment and maturation. Additionally, we identify candidate EPS proteins that may contribute to biofilm-associated physiology.

## Results

### Multi-omics profiling of early and mature stages of *M. smegmatis* biofilm provides insights into genes required for biofilm establishment and maintenance

In order to examine the genes that are crucial for the establishment of mycobacterial biofilm and its maintenance, we examined the transcriptomic and proteomic profile of the model organism, *Msm*, in its biofilm state. While transcriptomics provides an understanding regarding the regulatory mechanisms involved in biofilm formation, proteomics profiling enables functional characterization of the factors within the biofilm.

*Msm* is typically grown in a planktonic form in laboratory conditions. However, it forms biofilm under static culture conditions without detergents. In order to understand the genes responsible for biofilm formation, we profiled *Msm* transitioning from planktonic to biofilm (establishment) and its maintenance (maturation). *Msm* pellicular biofilm were grown in Sauton’s medium and samples were collected at two distinct stages. At 2-day post-inoculation, cells formed a thin layer at the air-media interface, which represents the establishment of biofilm and was designated as the initial stage (iBF). Following the 2nd day, a characteristic reticulation emerged, reaching its peak on day 5, which is regarded as the mature biofilm (mBF) (Fig. S1). Hence, day 2 and day 5 were selected for the analysis. We then employed RNA sequencing (RNA-seq) and label-free quantification based LC-MS to analyze the transcriptome and proteome profiles at each stage, respectively.

The transcriptome profiles of the bacteria exhibited significant differences, and a distinct separation was evident between the transcriptome profiles of planktonic, iBF, and mBF states (Fig. 1A and Table S1). Our data show that a transition from planktonic growth to initial biofilm formation involves a substantial reprogramming of the bacterial transcriptome. Compared to the planktonic state, we found that 730 genes exhibited a significant increase in transcript abundance (log_2_FC>1.5), whereas 332 genes displayed a decrease in transcript abundance (log_2_ FC<-1.5) during the initial (day 2) biofilm stage (Fig. 1B). A more pronounced shift in gene expression was observed when bacterial growth continued in the biofilm state (biofilm maturation; day 5). Compared to the initial stage, mature biofilm (mBF) displayed a significant increase in both upregulated (982 genes with log_2_ FC>1.5) and downregulated genes (1191 genes with log_2_ FC<-1.5) (Fig. 1C). This suggests a substantial reprogramming of the transcriptome as the biofilm matures. The most dramatic change in gene expression was observed between mature biofilms and planktonic cells. Compared to planktonic cells, mature biofilms displayed a significant upregulation of 1507 genes (log_2_ FC>1.5) and downregulation of 1433 genes (log_2_ FC<-1.5) (Fig. 1D). This extensive reprogramming highlights the importance of metabolic and physiological adaptations required by *Msm* to establish and thrive within the biofilm environment. Thus, a shift from planktonic to the biofilm state involves a dramatic reprogramming of gene expression, reflecting adaptation to the unique and complex biofilm microenvironment. Overall, the RNA-seq data revealed a regulation of gene expression taking place in two-phases during *Msm* biofilm establishment and maturation.

**Figure 1.**
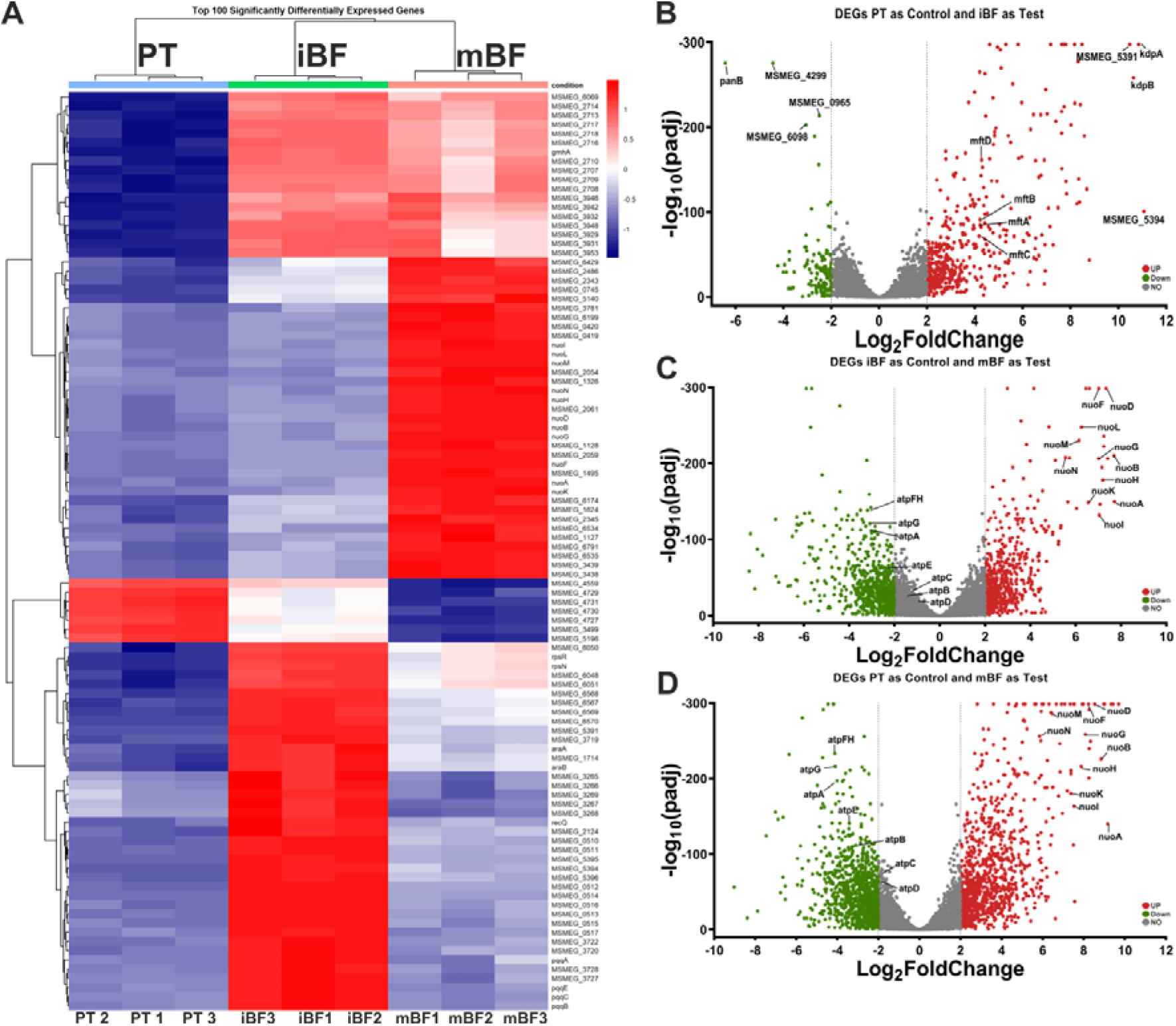
Differential expression of genes analyzed through transcriptomics analysis of the planktonic and biofilm states of *Mycobacterium smegmatis*. Shown here are the heatmap and volcano plots illustrating the differential expression of genes in *M. smegmatis* across different growth conditions. (A) Heatmap presenting the top 100 differentially expressed genes (DEGs) in *M. smegmatis* across three growth conditions: planktonic cells (PT), initial biofilm (iBF), and mature biofilm (mBF) stages. (B-D) DEGs between planktonic cells and the initial stage of biofilm, the initial stage of biofilm (iBF) and the mature stage of biofilm (mBF) and planktonic cells and the mature stage of biofilm (mBF), and, respectively. The x-axis in each case represents the fold-change on logarithmic scale, while the y-axis indicates the p-value as -log10(padj). Genes exhibiting a log_2_ fold change greater than +2 are coloured in red, whereas genes with a log2 fold change less than −2 are colored in green.

Similar to the transcriptomic analysis, our label-free quantification using high resolution nanoLC-MS approach revealed distinct transitions in proteome profiles during different stages of *Msm* biofilm formation (Fig. S2 and Table S2). In our experiments, we identified 33,568 unique peptides corresponding to 3819 proteins at a stringent cut-off of ≤1% false discovery rate (FDR). Among these proteins, 572 were differentially expressed in the initial biofilm (iBF), comprising 465 upregulated proteins (log_2_ FC>1.5) and 107 downregulated proteins (log_2_ FC<-1.5). The number of differentially expressed proteins increased in the matured biofilm (mBF) stage to 826 total proteins, with 663 upregulated and 163 downregulated compared to iBF. Furthermore, we observed that approximately 45 proteins were underrepresented in mBF relative to iBF, whereas nearly 118 proteins were overproduced during this maturation process. Notably, during the transition from iBF to mBF, 89% of proteins remained consistently upregulated and 87% remained downregulated.

These findings collectively provide insights into the intricate regulatory networks involved in mycobacterial biofilm formation and maturation, revealing critical genetic factors essential for the establishment and maintenance of biofilms in *Msm*. The integration of transcriptomic and proteomic data provides a holistic view of the molecular adaptations occurring during biofilm development. The circular representation of the expression data illustrates the dynamic interplay between the expression of the top genes and protein abundance across planktonic and initial as well as mature biofilm stages (Fig. 2A and 2C), highlighting critical pathways involved in biofilm establishment and maintenance. While the transcriptome and proteome show a fine correlation in iBF, there was very poor correlation in mBF (Fig. 2B and 2D). Importantly, transcriptomics primarily reflects the molecular drivers that initiate this transition; it might not be able to reflect the protein level, which is affected due to factors such as stability and accumulation of proteins. Moreover, biofilm mode of growth presents a matrix which may accumulate proteins, leading to poor correlation.

**Figure 2.**
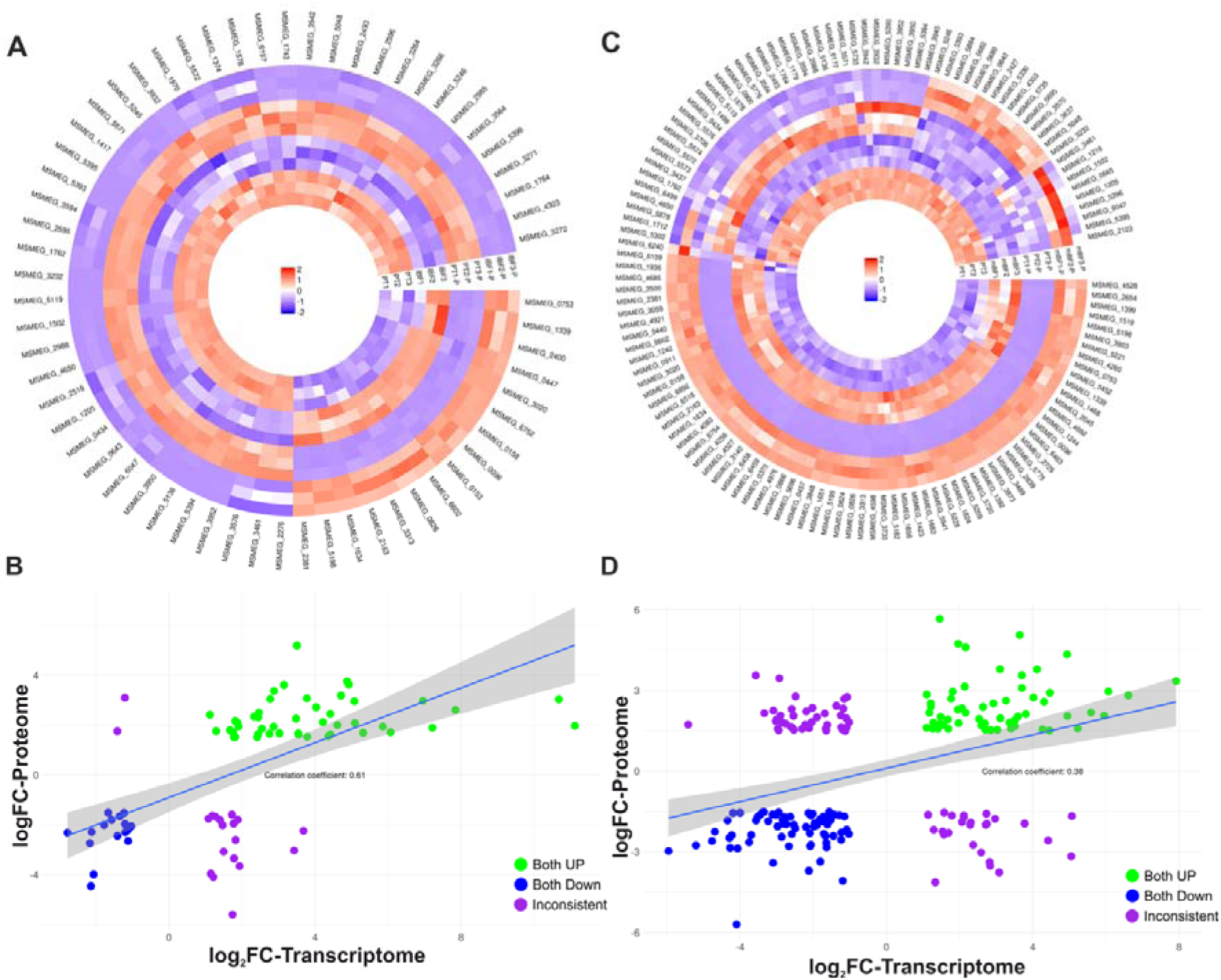
Comparison of the transcriptomic and proteomic profiles of planktonic and mature biofilm. (**A,C**) Heatmap of the relative abundance of the differentially expressed genes and proteins obtained from transcriptomic and proteomic profiles, respectively. (**A**) Planktonic (PT) vs initial (iBF) and (**C**) Planktonic (PT) vs mature biofilm (mBF) represent the replicates in transcriptome, while PT-P, iBF-P and mBF-P represent the replicates in the proteome. The gradient from blue to red indicates the relative degree of expression. The outer circle is the identity of each protein in *Msm.* (**B,D**) Correlation between the proteomic (y-axis) and transcriptomic (x-axis) DEGs obtained from the initial and mature biofilm state of *M. smegmatis* compared to planktonic growth, respectively. The blue and green points represent the down and upregulated genes, whereas purple points show inconsistency (profiles do not match).

### Extracellular polymeric substance of mycobacterial biofilm is composed of diverse proteins

Unlike other bacterial species such as *Staphylococcus aureus,* where specific EPS protein, extracellular adherence protein (Eap), can be used as marker for EPS quality (28), mycobacterial biofilms lack such readily identifiable proteins. Hence, acknowledging the limitations associated with EPS protein identification in mycobacterial biofilms is important. Nevertheless, in our experiments, the NaCl treatment of biofilm yielded a good amount of protein with visible bands on Coomassie-stained SDS-PAGE gels for the biofilm samples (Fig. S3). This is in contrast to barely visible protein bands in the planktonic controls. However, this observation also confirmed that NaCl did not by itself disrupt the cells, and that the proteins yielded while processing biofilm matrix are indeed matrix-associated proteins. To address the role of proteins, we performed proteomic analysis on EPS. We isolated the EPS from both initial and mature stages of biofilm using the NaCl treatment and subjected them to trypsin digestion and LC-MS/MS based identification.

Proteome analysis of the *Msm* biofilm EPS revealed 1106 proteins. These were mapped by 5597 unique peptides, of which 923 proteins were mapped with more than 2 unique peptides (Table S3). A volcano plot illustrating this proteomic data clearly highlights a significant number of proteins that are differentially regulated between the initial and mature stages of biofilm formation, revealing both increased and low representation of proteins in mature biofilms (Fig. 3A). Surprisingly, the localization profile of the EPS proteome revealed an overabundance of cytoplasmic proteins (∼29%) relative to membrane proteins (∼15%), and cell wall-associated proteins (∼3%) (Fig. 3B). The secretory proteins comprised only ∼4% of the total proteome as judged from the Uniprot assignment. Although we expected a higher proportion of secretory proteins, their low abundance could be a consequence of poor annotation of these specific proteins. Presence of cytoplasmic proteins in the EPS potentially emphasizes their moonlighting function in biofilm maintenance. Nevertheless, cell lysis, which is a well-documented phenomenon in biofilms,^20^ cannot be ruled out as a contributing factor for their presence.

**Figure 3.**
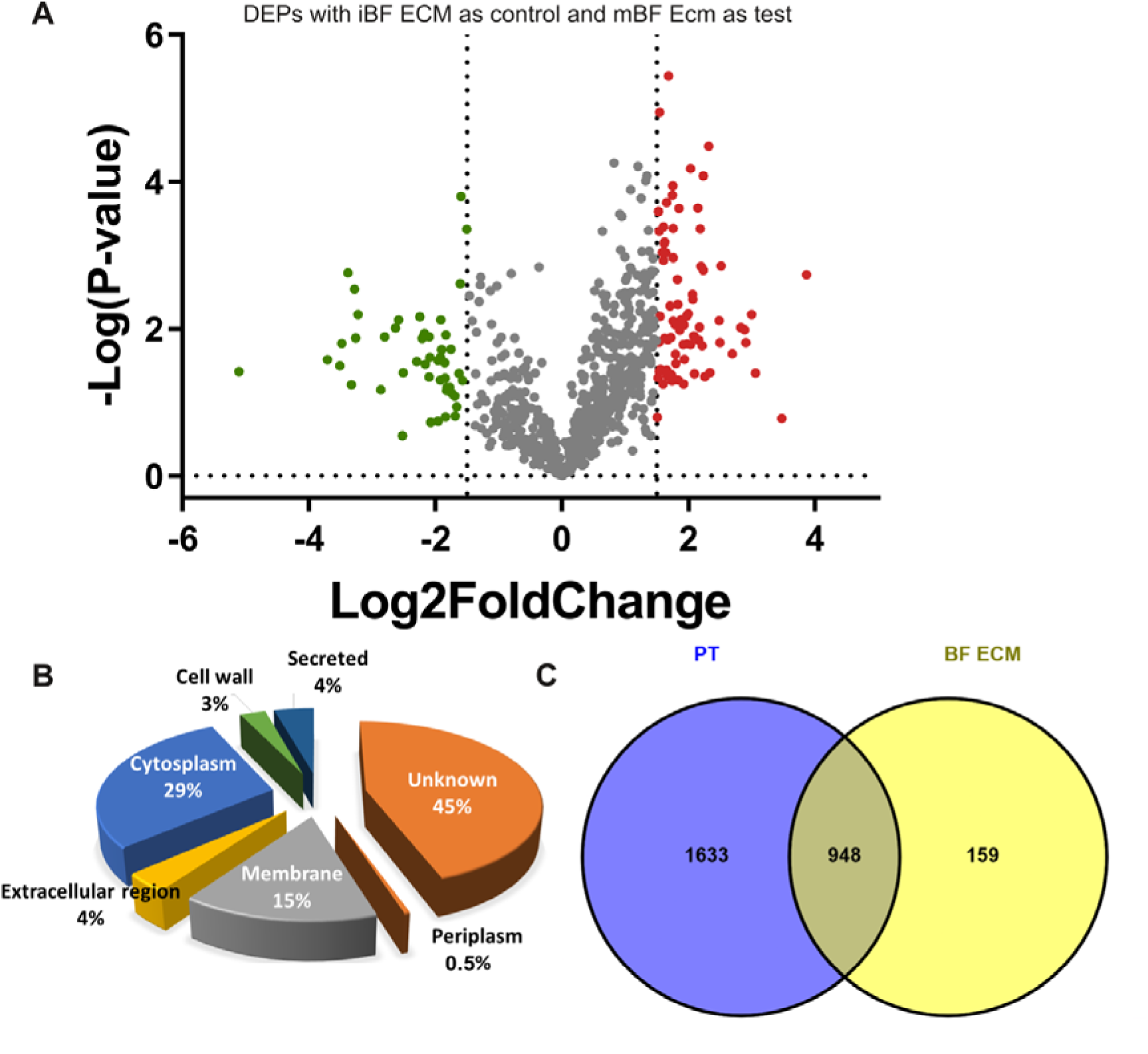
Proteomic profile and distribution of extracellular polymeric substance proteins in the initial and the mature stages of *M. smegmatis* biofilm. **(A)** Volcano plot illustrating the proteome profile of the extracellular polymeric substance (EPS) of the initial (iBF) and mature (mBF) stages of *M. smegmatis* biofilm. The x-axis represents the log_2_ fold change in protein abundance between the two stages, while the y-axis indicates the –log_10_ adjusted p-value, reflecting the statistical significance of the changes. Proteins that are significantly upregulated in the mature biofilm stage are indicated in red, whereas those downregulated are marked in green; proteins that show insignificant change are represented in grey. (**B**) A pie chart that illustrates the localization of all 1,106 EPS proteins identified in both the initial and mature stages of *M. smegmatis* biofilm. The chart visually represents the distribution of various EPS proteins across different cellular compartments or structures. Subcellular localization analysis was performed using data from UniProt. The protein distribution is shown for membrane, cell wall associated, cytoplasmic, periplasmic, secreted, extracellular region, and unknown as illustrated in UniProt. (**C**) Venn diagram comparing the proteins identified in the planktonic state of *M. smegmatis* with those identified at both the initial and mature stages of biofilm. The diagram highlights the overlap and unique proteins present in each growth condition.

Further, we employed a subtractive proteomic approach to identify proteins specifically associated with the biofilm EPS. We compared the planktonic state’s protein profiles, in which we identified 2581 proteins with the EPS fraction (initial and mature stages combined), isolated from *Msm* biofilms. This comparison identified a subset of 159 proteins enriched within the EPS (Fig. 3C), potentially representing key components or factors critical for biofilm formation and maintenance. Intriguingly, within this EPS-enriched fraction of 159 proteins, a noteworthy subset of 84 proteins either remains uncharacterized or has been categorized as hypothetical. Interestingly, 27 of these hypothetical proteins exhibit significant sequence conservation with proteins in *Mtb*, suggesting a similar pool of proteins involved in biofilm formation in *Mtb* as well.

The proteome analysis highlighted an array of proteins being differentially regulated between initial and mature stages of the biofilm. Notably, we identified ATP-dependent Clp proteases and PepD protease localized within the EPS of biofilms. Additionally, proteins showing an upregulation in the EPS were primarily nucleases, peptide synthetases, oxidoreductases, and conserved hypothetical proteins (Table S3). Very interestingly, BfrB, which is known to function as a nano-compartmental protein, was found to be significantly upregulated in the biofilm EPS. BfrB facilitates the storage of essential minerals such as iron within the bacterial cell.^21^ Transcriptomic analysis revealed a significant 3-fold upregulation of BfrB in both initial and mature biofilms as compared to planktonic cells. This upregulation was also corroborated with its presence in the biofilm EPS proteome. This suggests its likely secretion during biofilm development and highlights its potential and specific role in mycobacterial biofilm establishment and stability.

### Pathway enrichment analysis revealed enhanced oxidative stress response and metabolic adaptation in mature biofilm

Transcriptomic analysis revealed a pronounced shift in metabolic activity between planktonic cells and biofilm stages. In our study, we found a significant upregulation of oxidative stress-related pathways in the mature biofilm as compared to both planktonic cells and the initial biofilm stage. This was evidenced by the enrichment of genes involved in iron-sulphur cluster assembly (Fe-S and 4Fe-4S clusters), potassium uptake, and redox enzyme activities, including alcohol dehydrogenase, NADH dehydrogenase, nitrate reductase, and oxidoreductase (Fig. 4). These enzymes often function as key players in cellular redox balance and defense against oxidative stress.^22–26^ During oxidative stress and iron limitation Fe-S cluster proteins are degraded and bacteria respond by upregulating the *suf* genes in sulfur utilization pathway for reformation of the Fe-S clusters.^27^ Concurrently, a downregulation of ATP synthase activity, metallopeptidase, and hydrolase activities were observed, particularly in the mature biofilm stage (Fig. 4). This suggests a potential reduction in energy production and protein turnover, which might be a consequence of the increased metabolic burden imposed by oxidative stress. Interestingly, the upregulation of NADH dehydrogenase activity in the mature biofilm coupled with the downregulation of ATP synthase activity suggests a potential shift in energy metabolism. NADH dehydrogenase is a key component of the electron transport chain, typically linked to ATP production.^28^ However, the downregulation of ATP synthase implies that the generated reducing power (NADH) might not be fully utilized for ATP generation. This discrepancy could indicate a redirection of energy metabolism towards alternative pathways, such as biomass production or cellular maintenance processes to consume excessive reductive equivalent, or it might reflect an attempt to balance energy production with the increased demand for redox equivalents to counteract oxidative stress.

**Figure 4.**
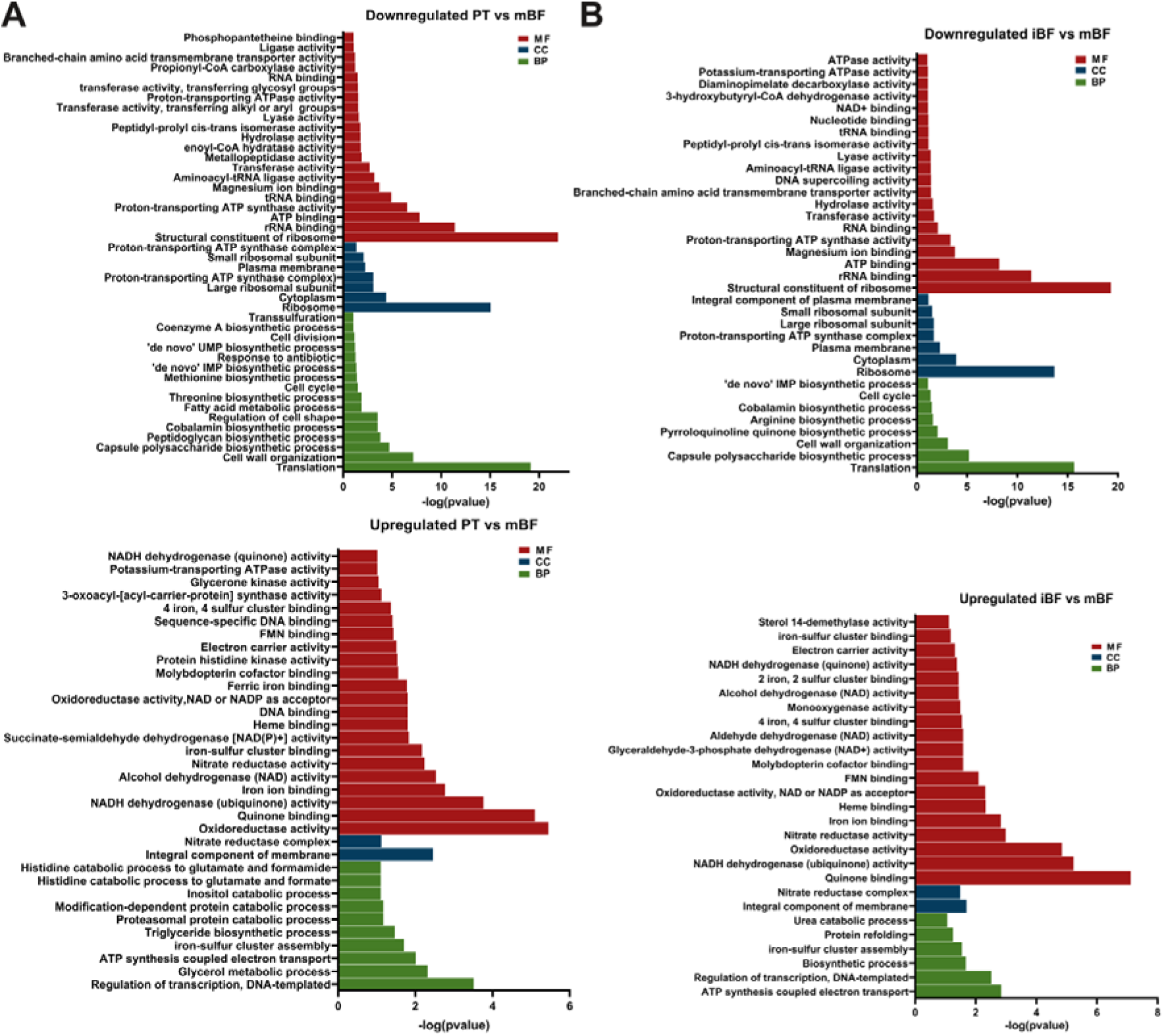
Dynamics of gene expression and pathway regulation in biofilms. The panels illustrate the dynamic changes in gene expression and pathway regulation during the transition from planktonic (PT) to mature biofilm (mBF), and initial stage (iBF) to mature stage of biofilm. The analysis was performed using the DAVID Gene Ontology (GO) tool, focusing on genes that exhibited expression changes >1.5 log2FC (upregulated) or <-1.5 log2FC (downregulated). y-axis in each case represents pathways identified through the GO analysis, whereas x-axis represents the -log(p-value) for each pathway, indicating the significance of the enrichment. Each bar in the panels corresponds to a specific biological pathway identified through the GO analysis. MF, molecular function; CC, cellular component; BP, biological process.

In contrast, the initial biofilm stage exhibited a more moderate response to oxidative stress compared to the mature biofilm (Fig. S4). While upregulation of transmembrane transporter activity, potassium-transporting ATPase, iron-sulphur cluster binding proteins, and redox enzymes was apparent, it was less pronounced than in the mature biofilm. These findings suggest that the initial biofilm is experiencing an adaptive response to oxidative stress, but a thorough response is yet to be established. These results collectively indicate that the mature biofilm environment is characterized by elevated oxidative stress. The upregulation of oxidative stress response pathways suggests that biofilm cells have activated defense mechanisms to counteract the detrimental effects of reactive oxygen species. The downregulation of energy-related processes, including ATP synthase, coupled with the upregulation of NADH dehydrogenase, points to a potential metabolic shift in the mature biofilm, likely in response to the increased oxidative stress.

### Mycobacterial biofilm establishment and maintenance involves substantial reprogramming of energy metabolism

We shortlisted the top 100 genes showing the highest level of expression changes between different stages of biofilm cycle (Fig. 1A). Interestingly, the *nuo* operon, encoding NADH dehydrogenase I, was significantly upregulated in the mature biofilm state (mBF) as compared to both the initial biofilm state (iBF) and the planktonic culture (Fig. 1A). The *nuo* operon is primarily involved in the NADH/NAD+ metabolism. Interestingly, all 14 genes within the *nuo* operon (*nuoA-N*) along with its regulatory gene, MSMEG_2064, displayed a nearly 7-fold increase in expression in the mature biofilm as compared to the initial biofilm state (Table S1). MSMEG_2064 was also expressed 2-fold as compared to the planktonic state, suggesting a strong activation of the *nuo* operon during biofilm maturation. It is important to note that *ndh*, a single-copy gene encoding NADH dehydrogenase II, plays a similar role as the *nuo* operon genes. However, unlike the *nuo* operon, *ndh* expression did not show any significant difference between the initial biofilm stage, mature biofilm stage, and planktonic cells (Table S1). Since the *nuo* operon showed an overexpression in the mature stage of biofilm cycle, we suspected a shift in NADH/NAD+ metabolism also. Our NADH/NAD^+^ estimation using *in vivo* ratiometric sensor peredox-mCherry showed a continuous increment in the NADH levels while shifting from planktonic stage to initial stage to mature stage of biofilm (Fig. 5A). The NADH levels were saturated and showed no increase after day 4 when the mature stage is formed.

**Figure 5.**
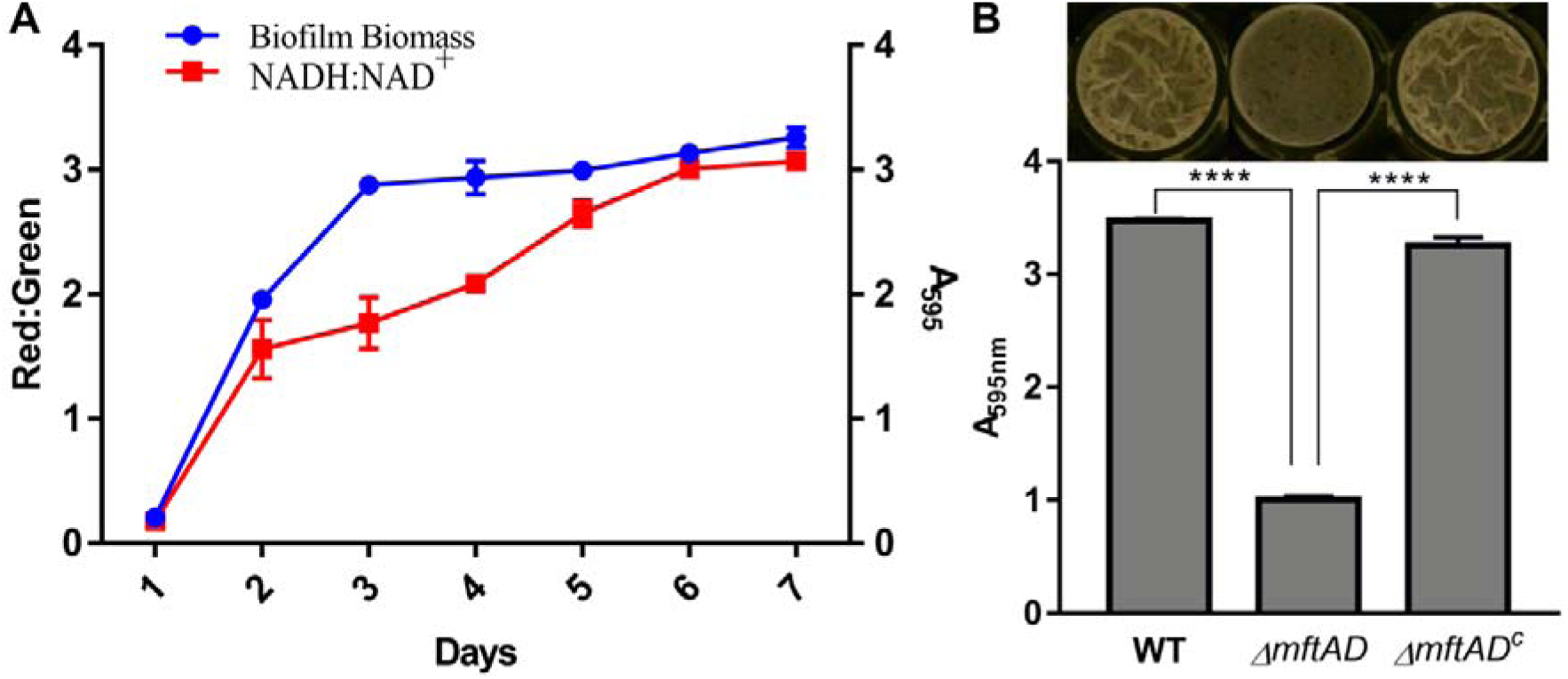
Role of redox balance machinery and electron acceptors in biofilm establishment. (**A)** Dynamic relationship between the intracellular NAD^+^:NADH ratio and biofilm formation in *M. smegmatis*. The NAD^+^:NADH ratio, an indicator of cellular redox state, was determined using the pMV762-Peredox-mCherry plasmid. Biofilm biomass accumulation was quantified using the crystal violet staining method. NAD^+^:NADH ratio (Red:Green fluorescence ratio) is shown on the right, whereas biofilm quantification as absorbance at 595 nm is shown on the left. The data are shown from day 1 to day 7. (**B**) 5-day-old biofilm formed at the liquid-air interface in the presence of glucose as the carbon source by the three bacterial strains viz. WT, Δ*mftAD* (mycofactocin biosynthesis operon-deleted strain) and Δ*mftAD^c^* (complemented strain). The graph below shows the quantification of the biofilm using the crystal violet dye assay. The absorbance measured at 595Lnm in each case is plotted. In both the panels, the experiments were carried out least thrice, and the data represent an average of these experiments. In panel B, only one representative image of biofilm formation is shown on the top. ‘****’, P-value□<□0.0001.

In contrast to the *nuo* operon, the ATP synthase complex consisting of eight genes (*atpBEFHAGDC*) showed a 4-fold downregulation during the mature stage (mBF) as compared to planktonic cells (Table S1). Interestingly, no significant difference in expression was observed between the initial stage (iBF) and planktonic cells for ATP synthase complex. This finding strongly suggests a substantial shift in energy metabolism within the mature biofilm. Our data thus demonstrate a dynamic shift in gene expression during mycobacterial biofilm development. The upregulation of the *nuo* operon for NADH production in mature biofilms and the downregulation of the ATP synthase complex indicate a potential metabolic reprogramming strategy.

### Mycofactocin is essential for the mycobacterial biofilm establishment

Our transcriptomics analysis revealed a 4-fold increase of alcohol dehydrogenase genes and the *mftABCDEF* gene cluster involved in the biosynthesis of mycofactocin, an electron acceptor for these enzymes, in the initial stages of biofilm formation and reduced to planktonic levels in mature biofilms (Table S1).^29,30^ This finding suggests a potential role for mycofactocin dependent energy metabolism during biofilm establishment. We, therefore, expanded our investigation by studying the effect of *mftAD* knockout on biofilm formation. The *mftAD* knockout failed to develop mature biofilms (Fig. 5B), whereas the complemented strain showed restoration of biofilm formation (Fig. 5B). This indicated the specificity of *mftAD* in the maturation of biofilm beyond the initial stage and suggests a potential role of mycofactocin and the alcohol assimilation pathway in mycobacterial biofilm establishment.

## Discussion

Biofilm formation is a complex process that requires a dynamic interplay between gene expression and protein localization to build the multicellular structure embedded in an extracellular polymeric substance (EPS) made of DNA, lipids and protein. Growth in this protective environment is also responsible for the emergence of drug resistance in pathogenic bacteria. Prolonged chemotherapy in mycobacterial diseases correlates its ability to grow in biofilms on human host.^31^ Previous studies have identified the presence of mycolic acid,^8^ cellulose,^31^ and extracellular DNA (eDNA)^32^ in the EPS of mycobacterial biofilms. However, the role of proteins in biofilm integrity has remained largely unexplored. Interestingly, the complete disruption of mycobacterial biofilms upon treatment with proteinase K underscores the critical importance of proteins in the EPS.^31^ Nevertheless, a comprehensive understanding of the specific proteins present in the mycobacterial biofilm EPS is currently lacking. While transcriptomic analysis provides valuable insights into gene expression, it does not directly translate to protein function within the biofilm’s EPS. NaCl extract of the EPS revealed an unexpected protein distribution. Contrary to our initial expectation of a predominance of secreted proteins, we observed that cytoplasmic proteins constitute the most abundant fraction within the EPS, followed by membrane and cell wall proteins (Fig. 4). This finding challenges the traditional paradigm of biofilm EPS composition and suggests potential moonlighting functions for cytoplasmic proteins within this environment. These proteins are either actively transported to the EPS or released from lysed cells during biofilm development. We hypothesize that these proteins are required for structural integrity, nutrient acquisition, and other biofilm-associated processes. The low abundance of secretory proteins observed in our analysis could also be attributed to limitations in current protein annotation for these specific molecules.

Interestingly, our data identified Clp proteases, ClpP1 and ClpP2, within the EPS of both initial and mature biofilms. Clp proteases are essential for protein quality control and play a critical role in bacterial survival.^33^ Their presence in the EPS suggests a potential role in regulating protein turnover within this environment. Importantly, studies in other bacterial species like *Bacillus subtilis* have shown that Clp proteases can become ATP-independent in the presence of ADEP-4 antibiotic.^34^ This finding highlights the potential of Clp proteases as attractive targets for novel drugs aimed at disrupting protein homeostasis and compromising biofilm integrity. Furthermore, our data highlight the differential regulation of EPS proteins between the initial and mature stages of the biofilm. Abundance of specific protein groups, including nucleases, peptide synthetases, oxidoreductases, and conserved hypothetical proteins (Table S3), suggests dynamic changes in the EPS composition throughout biofilm development.

Our transcriptomic and proteomic analyses revealed a compelling link between iron homeostasis and *Msm* biofilms. We observed a significant (3-fold) upregulation of the bacterioferritin protein (BfrB) during biofilm development and within the EPS proteome. The presence of BfrB in the biofilm EPS suggests a previously unexplored aspect of iron regulation in *Msm* biofilms.^9,35,36^ More investigation of BfrB’s role in iron homeostasis, its potential interaction with Mt-Enc homologs,^21^ and its impact on biofilm development will offer valuable insights into the mechanisms that govern biofilm formation in mycobacteria.

This study highlights the unique metabolic rewiring that the mycobacteria undertake within biofilms. Our findings suggest that mycobacteria preferentially rely on NADH/NAD+ metabolism for energy generation during biofilm establishment and maintenance. Excess reducing equivalents is plausibly generated to counter the oxidative stress during the initial stages of biofilm formation. This shift in metabolic dependency strongly suggests that targeting NADH/NAD+ metabolism would be beneficial in curtailing infections involving mycobacterial biofilm, which has been shown to be present in the animal lung during tuberculosis infection.^37–39^ Compounds such as the 2-mercaptobenzothiazoles and 2-metrcaptoquinazolinones inhibiting the type II NADH dehydrogenases will, therefore, prove efficacious in eradicating persistent mycobacterial infection.^40,41^ Additionally, our observation aligns with another interesting finding that downregulation of the ATP synthase complex in bacteria can lead to an enhanced proton motive force (PMF). The PMF is a critical factor in various cellular processes, including nutrient uptake and efflux. By potentially downregulating ATP synthase while maintaining a robust PMF, mycobacteria within biofilms can optimize their energy expenditure for survival and persistence.^42–44^ We also observed that mycofactocin, a factor required for high NADH associated methanol and ethanol metabolism, plays a significant role in biofilm formation and maturation. This suggests that mycofactocin may contribute to the metabolic adaptations necessary for the persistence of mycobacteria within biofilms, particularly under conditions of metabolic stress.

In conclusion, this study provides a comprehensive view of the genetic and metabolic adaptations underlying mycobacterial biofilm development. Through integrated omics analyses, it reveals that biofilm development is a highly dynamic, two-phase process characterized by extensive reprogramming of gene expression, protein abundance, and metabolic pathways. The findings highlight a pronounced shift in energy metabolism, marked by increased NADH generation and reduced ATP synthesis, alongside enhanced oxidative stress responses that support survival in the biofilm environment. The identification of diverse EPS associated proteins underscores the structural and functional complexity of the biofilm matrix. Collectively, these results advance our understanding of mycobacterial biofilms and identify potential molecular targets for disrupting biofilm-associated persistence.

## Materials and Methods

### Bacterial growth and biofilm quantification

Wild-type *M. smegmatis* mc^2^155 and its transformed variants were cultivated in Difco’s Middlebrook 7H9 (MB7H9) broth, supplemented with 2% glucose and 0.05% Tween 80 and incubated at 37°C with continuous agitation at 200 rpm. Bacterial growth was evaluated by periodically measuring the optical density of the culture at 600 nm (OD600) and constructing corresponding growth curves.

Biofilm experiments were conducted by initially culturing all strains in MB7H9 medium supplemented with 2% glucose and 0.05% Tween 80 at 37°C until they reached the stationary phase (∼72 h). Subsequently, we inoculated secondary cultures in MB7H9 medium at OD600 ∼ 0.6, and used the log-phase cultures (∼12 h) to initiate biofilm formation. To facilitate biofilm formation, the bacterial strains were grown undisturbed in Sauton’s medium (HiMedia) in a 24-well plate, supplemented with 2% glycerol and 2% acetamide, following established methods.^45^ The plates were then sealed with parafilm and incubated at 37°C in a humidified chamber for varying number of days as required. For biofilm quantification, we modified a previously described protocol.^46^ Briefly, we emptied and dried the wells containing the biofilm, stained them with 1% crystal violet, rinsed them with water, and allowed them to air dry. The bound dye was subsequently dissolved in acetic acid, and the resulting eluate was analyzed by measuring the absorbance at 595Lnm.

### Biofilm EPS extraction

Biofilm EPS extraction was performed as described elsewhere.^47^ Briefly, biofilms that were grown in 24 well plates were collected and resuspended in different concentrations of NaCl. Before the centrifugation, they were incubated for 10 min at room temperature. The suspensions were centrifuged at 5000 rpm for 10 min at room temperature and the supernatants were transferred into new vials as NaCl fractions, and further processed for protein isolation.

### Measurement of NADH/NAD+ ratio

*M. smegmatis* cultures containing pMV762-peredox-mCherry^39^ were inoculated into 5 ml of 7H9 MB media supplemented with 2% glucose, 0.05% Tween-80, and 50 μg/ml hygromycin. The cultures were grown to mid-log phase (OD600 ∼ 0.8) and used to initiate biofilm formation. Biofilms were harvested from days 1 to 7 and washed with PBS. A 200 µl aliquot of each biofilm sample was loaded into a 96-well plate, and spectra were recorded using a hybrid fluorimeter with an absorption wavelength of 400 nm and 587 nm, while emission was measured at 615 nm and 510 nm. The fluorescence emission ratio at 615 nm to 510 nm (red/green) was plotted against time to assess the NADH/NAD^+^ ratio within the bacterial cells.

### RNA extraction and transcriptomic analysis

RNA was extracted from bacterial cultures at different growth stages, including log phase culture grown till 0.6 OD, initial biofilm, 2-day-old biofilm, and mature biofilm (5 days old), using the Qiagen mini kit by following manufacturer’s instructions. Quality control was performed by quantitating RNA samples on a Qubit fluorimeter and determining RNA integrity on an Agilent Tapestation 4200. Subsequently, 1 μg of RNA samples underwent bacterial rRNA depletion using the NEBNext rRNA depletion kit, followed by DNase I digestion and purification using RNA purification beads. Double-stranded cDNA libraries were constructed from the rRNA-depleted RNA using the NEBNext Ultra II Directional RNA Library prep kit for Illumina, including RNA fragmentation, first strand cDNA synthesis, second strand cDNA synthesis, end repair, adapter ligation, and PCR enrichment. Cleaned libraries were assessed for size range and average library size. Quality of reads was evaluated with FastQC and trimmed using Fastp. Transcriptome analysis was conducted by mapping reads onto the reference transcriptome using bowtie2 and quantified with featureCounts. Removal of highly abundant rRNA, tRNA, and sRNA counts was performed to avoid confounding effects. Differential expression analysis was conducted using DESeq2, with genes filtered based on false discovery rate (padj) < 0.05 and minimum expression log_2_ fold change (FC) ≥ 2.

### Whole cell Protein purification and LC-MS/MS analysis

Proteins isolated from different stages of the biofilm and from the extracellular polymeric substance (EPS) were precipitated using Trichloroacetic acid (TCA) in a 1:4 (v/v) ratio. Protein pellets then solubilized in 6M urea and quantified with BCA (Sigma). 10 μg of proteins were reduced and alkylated using DTT and iodoacetamide, respectively. These then Trypsinised at pH 7.8 in ammonium bicarbonate at 37°C overnight. Desalting was done using custom made EmporeTM C-18 stage tips conditioned with 100% acetonitrile and equilibrated with 0.1% formic acid solution. Samples were loaded and washed with 0.1% formic acid and subsequently eluted in 50% acetonitrile with 0.1% formic acid. Proteins were then vacuum dried and re-suspended in 0.1% formic acid. 600 ng of peptides were separated on RSLC nano-UPLC using C-18 column, 2 μm particle size, 100 Å pore size, 75 μm × 50 cm (PepMapTM RSLC). Solvents used were 0.1% formic acidic in water (solvent A) and 0.1% formic acidic in acetonitrile (solvent B). Separation was achieved with gradient increase in solvent B from 5% to 25% in 30 min. MS spectra of separated peptides were obtained using Orbitrap at 60,000 resolution equipped with nanospray source (Thermo Fisher Scientific). The spray ionization was performed at 1.8 kV voltage and 275°C capillary temperature. MS data was acquired in full scan mode from 350 to 2000 m/z range. MS2 was performed on the 20 most abundant MS1 parent ions. The fragmentation was performed using collision induced dissociation at normalized collision energy 35 eV. Peptide identification was performed suing MaxQuant 3.0, identification was performed using reference proteome of *Mycobacterium smegmatis* mc^2^155 version 5 (https://mycobrowser.epfl.ch/). Following parameters were used for identification of the proteins: Enzyme specificity, Trypsin/P; maximum missed cleavages, 2; mass tolerance for first and main search, 20 and 4.5 ppm respectively; mass tolerance for fragment ion, 0.5 Da; fixed modification, carbamidomethylation (C); variable modifications used were N-terminal acetylation and methionine oxidation; minimum unique peptide required for identification, 1; minimum peptide length for identification, 7; max. peptide mass, 4600 Da; PSM identification and protein inference FDR were set at 0.01; dependent peptide and match between run was enabled. Protein group files generated by Maxquant were further used for the differential expression analysis using Perseus. After removal of contaminations, reverse sequences and modified peptides proteins with valid label free quantification (LFQ) intensities in all three replicates of all conditions were selected for calculating Log_2_FC and significance was calculated using t-test.

## Acknowledgements

HN and RS thank Department of Biotechnology, Govt. of India for the fellowship. The work is supported by a grant (#27(0325)/17/EMR-II) from the Council of Scientific and Industrial Research, Govt. of India to VJ, and by the intramural funds from IISER Tirupati to RM. We thank Gokul Nair for proofreading the manuscript and suggestions. We also thank Lokdeep Teekas for help with analysing transcriptomics and proteomics data.

## Competing interests

All authors declare no financial or non-financial competing interests.

## Data availability

All data generated or analyzed during this study are included in this article and its supplementary information files.

## Supplementary figure legends

**Figure S1**. **Development of *M. smegmatis* biofilm**. Formation of *M. smegmatis* biofilm with time is shown here. *M. smegmatis* biofilm was observed and imaged over a period of 7 days as mentioned in a 24-well plate. One well in each case is imaged and shown. The experiments were repeated multiple times; only one representative image in each case is shown here.

**Figure S2. Differential expression of proteins observed through proteomics analysis of the planktonic and biofilm states of *Mycobacterium smegmatis*.** Shown here are the volcano plots and heatmap illustrating the differential expression of proteins in *M. smegmatis* across different growth conditions. Panels A, B, and C show the differentially expressed proteins (DEPs) between planktonic cells and the initial stage of biofilm (panel A), planktonic cells and the mature stage of biofilm (panel B), and initial and mature stages of biofilm (panel C). The x-axis in each case represents the fold-change on logarithmic scale, while the y-axis indicates the p-value as -log10(padj). Proteins with a -log10(padj) value greater than 1.5 are highlighted in red, while those with a value less than 1.5 are shown in black. Panel D shows heatmap representing the top 100 DEPs in *M. smegmatis* across three growth conditions in replicates: Planktonic cells (PT), initial biofilm (iBF), and mature biofilm (mBF) stages.

**Figure S3. Examination of biofilm EPS proteins using sodium chloride method.** EPS proteins present in the *M. smegmatis* biofilm were extracted using NaCl method as described in the materials and methods, and were examined on SDS-PAGE gel. Shown here are the images of Coomassie-stained SDS-PAGE gel of extracted EPS proteins from Biofilm (A). Different concentrations of NaCl as given were used for extraction. Planktonic culture (B) was used as negative control. In both panels, ‘L’ represents the molecular weight marker with few bands marked.

**Figure S4. Dynamics of gene expression and pathway regulation in the biofilm.** The panels illustrate the dynamic changes in gene expression and pathway regulation during the transition from planktonic (PT) to initial stage (iBF) of biofilm. The analysis was performed using the DAVID Gene Ontology (GO) tool, focusing on genes that exhibited expression changes >1.5 log_2_FC (upregulated) or <-1.5 log_2_FC (downregulated). y-axis in each case represents pathways identified through the GO analysis, whereas x-axis represents the -log(p-value) for each pathway, indicating the significance of the enrichment. Each bar in the panels corresponds to a specific biological pathway identified through the GO analysis. MF, molecular function; CC, cellular component; BP, biological process.

## Supplementary Tables

Table S1: Transcriptome profiles of the planktonic, initial, and mature biofilm states.

Table S2: Proteomics profiles of the planktonic, and the extracellular polymeric substance of the initial and mature biofilm.

Table S3: Mapping of the EPS proteins identified using LC-MS/MS mass spectrometry.

## Notes

### Competing Interest Statement

The authors have declared no competing interest.

