## Supplementary figures for "From planktonic to sedentary lifestyle: Molecular dissection of the establishment and maintenance of mycobacterial biofilm"

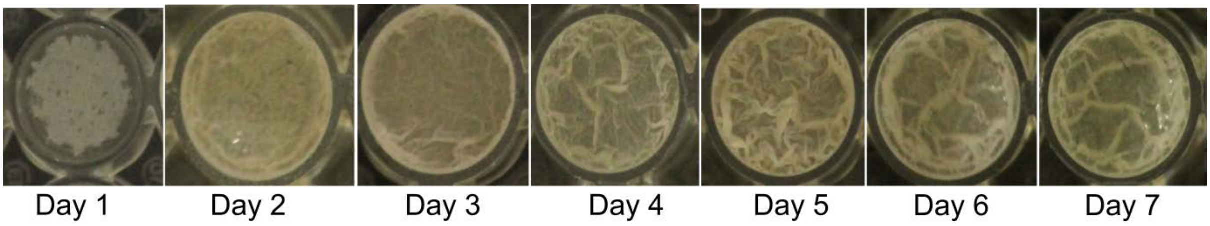


**Figure S1**. **Development of *M. smegmatis* biofilm**. Formation of *M. smegmatis* biofilm with time is shown here. *M. smegmatis* biofilm was observed and imaged over a period of 7 days as mentioned in a 24-well plate. One well in each case is imaged and shown. The experiments were repeated multiple times; only one representative image in each case is shown here.

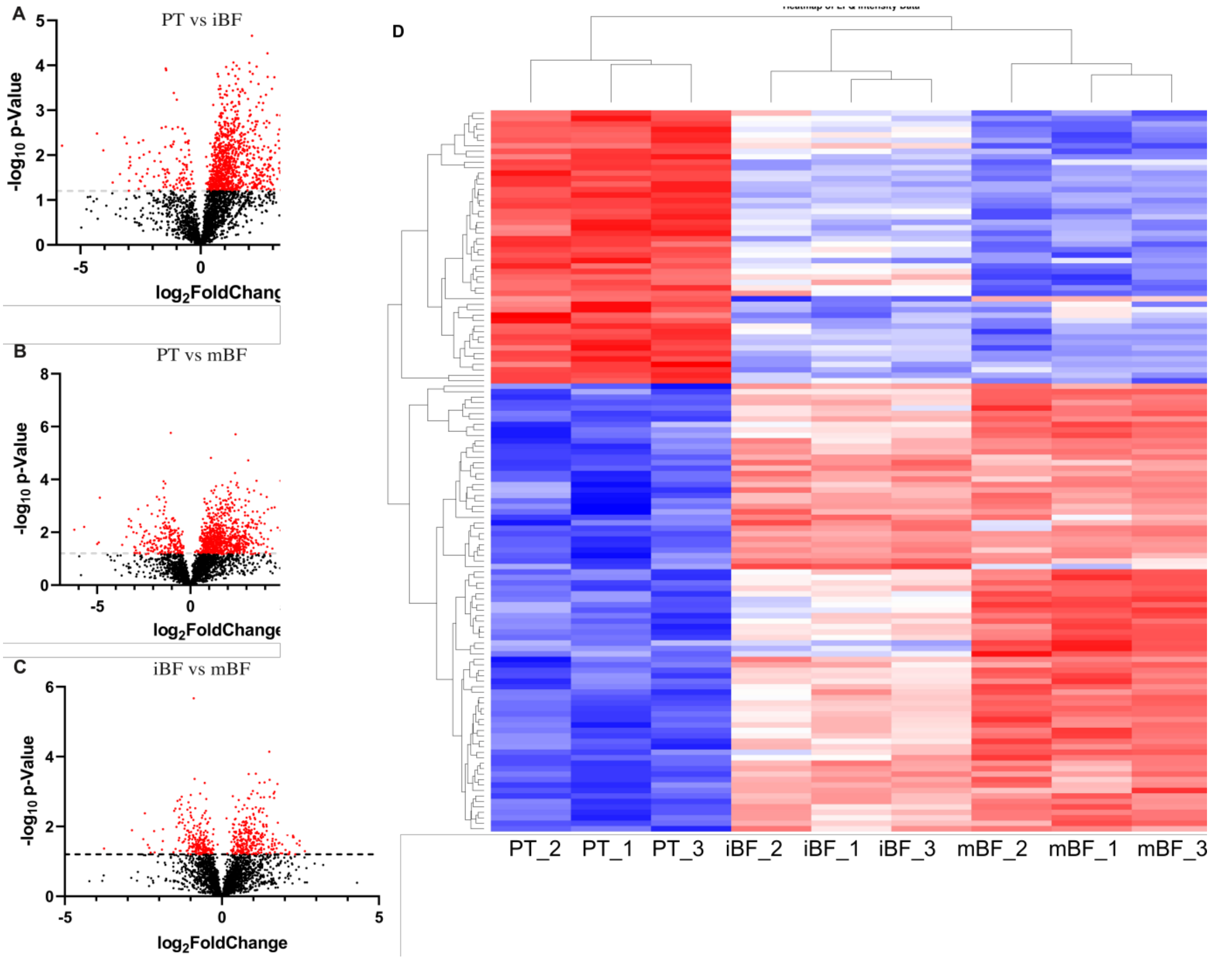


**Figure S2. Differential expression of proteins observed through proteomics analysis of the planktonic and biofilm states of *Mycobacterium smegmatis*.** Shown here are the volcano plots and heatmap illustrating the differential expression of proteins in *M. smegmatis* across different growth conditions. Panels A, B, and C show the differentially expressed proteins (DEPs) between planktonic cells and the initial stage of biofilm (panel A), planktonic cells and the mature stage of biofilm (panel B), and initial and mature stages of biofilm (panel C). The x-axis in each case represents the fold-change on logarithmic scale, while the y-axis indicates the p-value as -log10(padj). Proteins with a -log10(padj) value greater than 1.5 are highlighted in red, while those with a value less than 1.5 are shown in black. Panel D shows heatmap representing the top 100 DEPs in *M. smegmatis* across three growth conditions in replicates: Planktonic cells (PT), initial biofilm (iBF), and mature biofilm (mBF) stages.

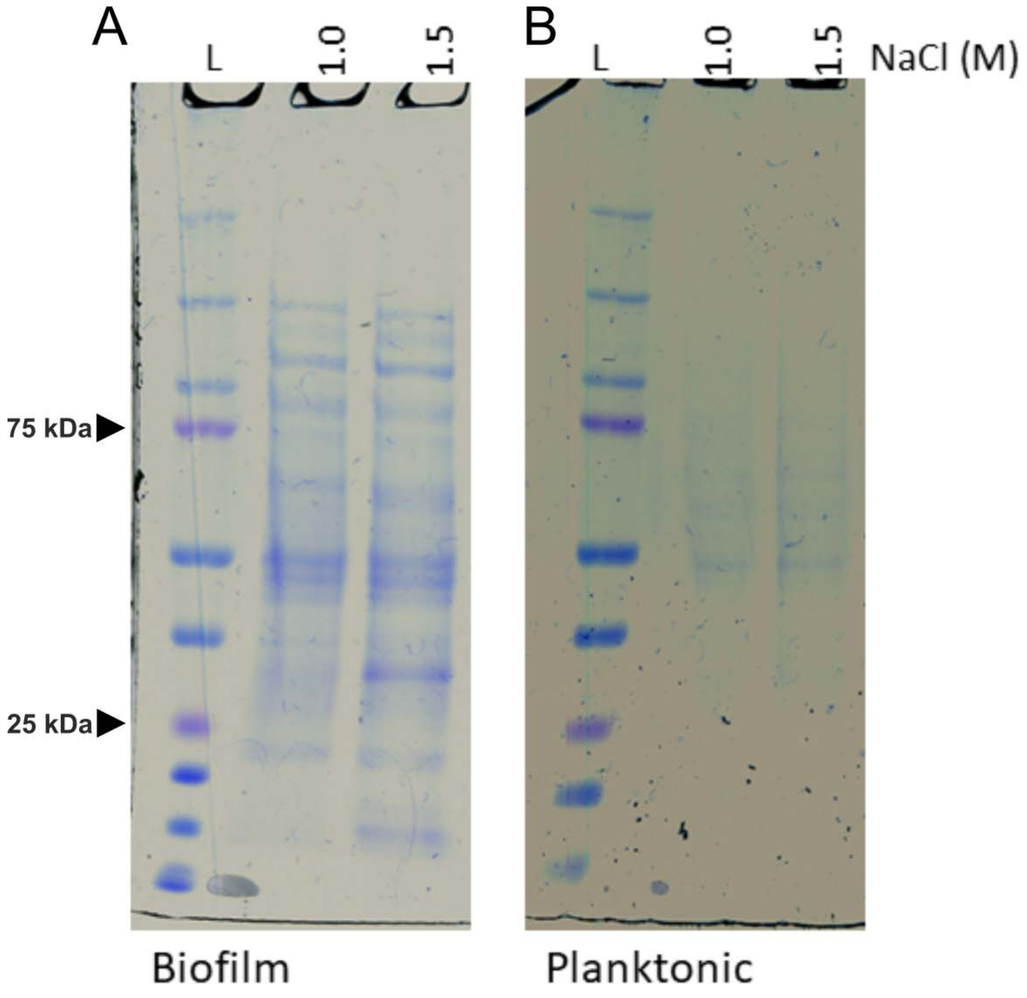


**Figure S3. Examination of biofilm EPS proteins using sodium chloride method.** EPS proteins present in the *M. smegmatis* biofilm were extracted using NaCl method as described in the materials and methods, and were examined on SDS-PAGE gel. Shown here are the images of Coomassie-stained SDS-PAGE gel of extracted EPS proteins from Biofilm (A). Different concentrations of NaCl as given were used for extraction. Planktonic culture (B) was used as negative control. In both panels, 'L' represents the molecular weight marker with few bands marked.

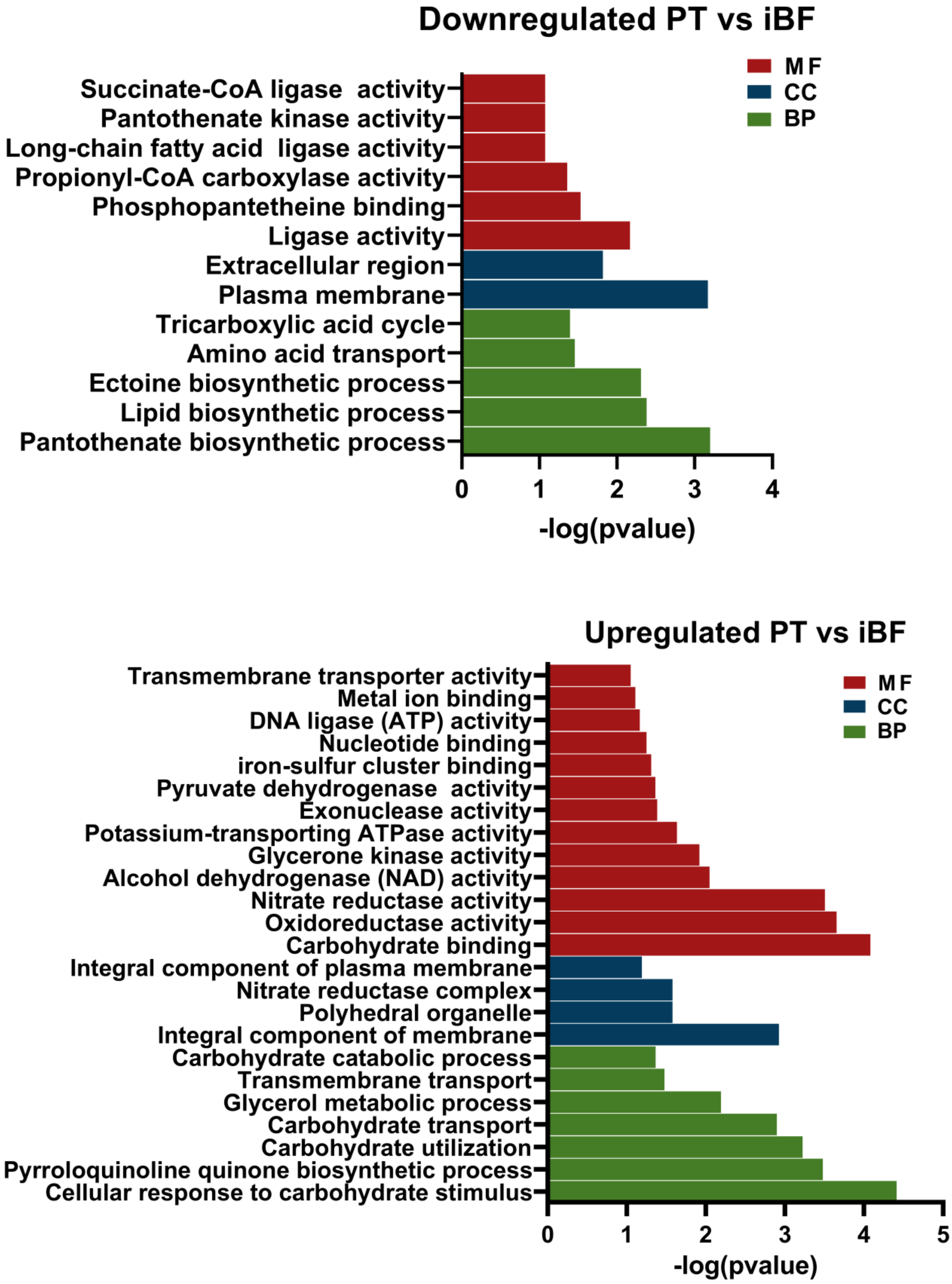


**Figure S4. Dynamics of gene expression and pathway regulation in the biofilm.** The panels illustrate the dynamic changes in gene expression and pathway regulation during the transition from planktonic (PT) to initial stage (iBF) of biofilm. The analysis was performed using the DAVID Gene Ontology (GO) tool, focusing on genes that exhibited expression changes >1.5 $log_2$FC (upregulated) or <-1.5 $log_2$FC (downregulated). y-axis in each case represents pathways identified through the GO analysis, whereas x-axis represents the -log(p-value) for each pathway, indicating the significance of the enrichment. Each bar in the panels corresponds to a specific biological pathway identified through the GO analysis. MF, molecular function; CC, cellular component; BP, biological process.
